# Polysaccharide BAP1 of *Bifidobacterium adolescentis* CCDM 368 attenuates ovalbumin-induced allergy through inhibition of Th2 immunity in mice

**DOI:** 10.1101/2024.09.14.613063

**Authors:** Katarzyna Pacyga-Prus, Tereza Hornikova, Dagmar Šrůtková, Katarzyna Leszczyńska-Nowak, Agnieszka Zabłocka, Martin Schwarzer, Sabina Górska

## Abstract

Allergies have become a growing problem and the number of cases is increasing yearly. Administration of postbiotics, well-defined bacterial molecules, is gaining attention as a novel and promising strategy to ameliorate the allergic burden. The BAP1 polysaccharide (PS) of *Bifidobacterium adolescentis* CCDM 368, was previously characterized by us regarding its structure and *in vitro* immunomodulatory properties. Here, to decipher the effect of BAP1 on immune system development, it was intranasally (i.n.) administered to germ-free mice. We observed increased IgA in bronchoalveolar lavage (BAL) fluid, decreased CCL2 production, and higher *Rorc* gene expression in the lung. The intranasal administration of BAP1 reduced lung inflammation and decreased eosinophils numbers in BAL in the ovalbumin-induced allergy mouse model. Moreover, BAP1 decreased OVA-specific IgE levels in sera and Th2-related cytokines in OVA-stimulated splenocytes and lung cells. Finally, increased *Rorc* and inhibited *Il10* gene expression were observed in lung tissue indicating their possible role in BAP1 function. Our findings support and expand on our previous *in vitro and ex vivo* studies by demonstrating that BAP1, with a unique chemical structure, induces a specific immunomodulatory effect in the host and could be potentially used for alleviating allergic diseases.

**Graphical abstract:** 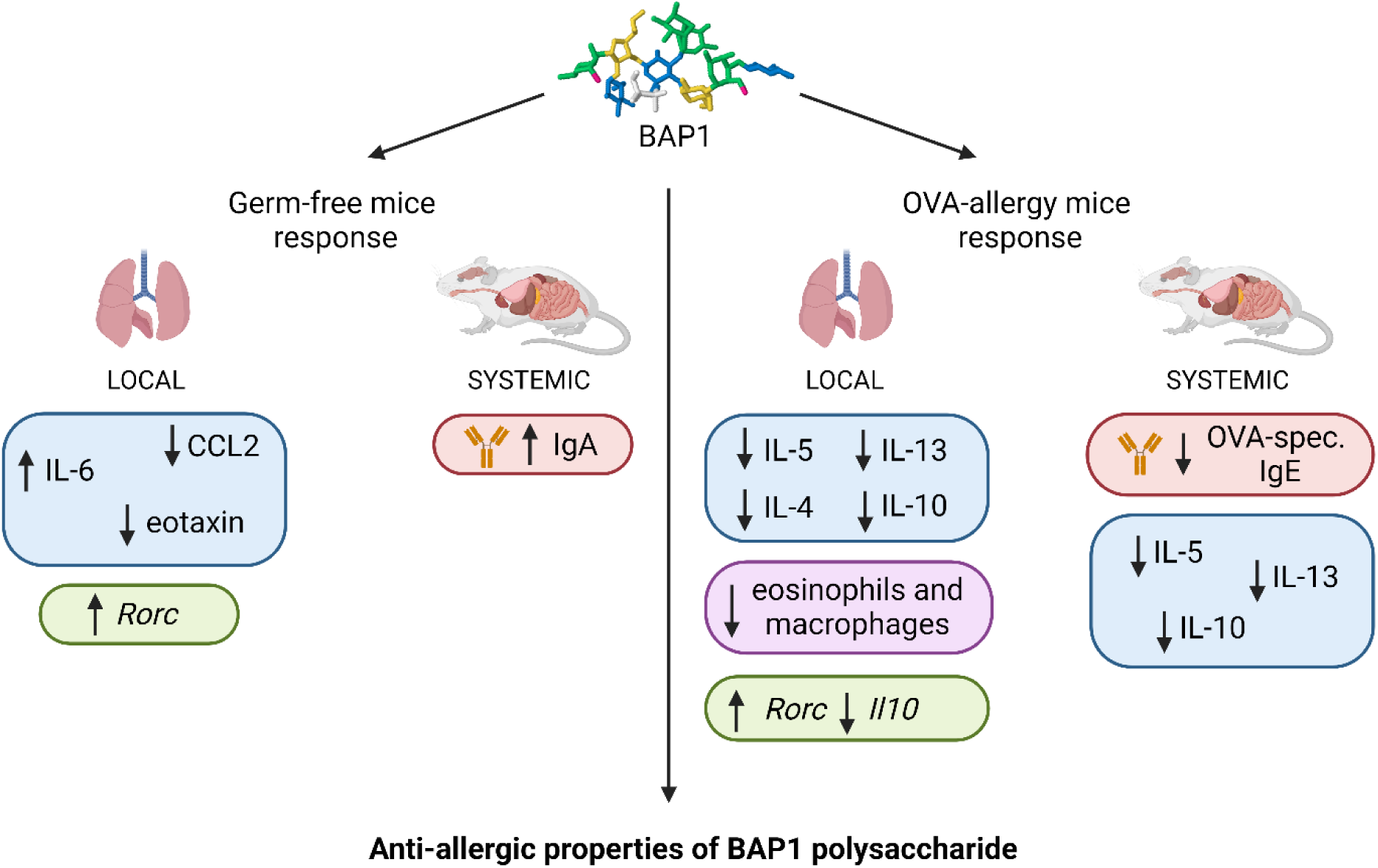

## 1. Introduction

Airway allergies have become a major healthcare problem affecting people worldwide. According to data collected by the European Academy of Allergy and Clinical Immunology (EAACI), in 2019 up to 40% of the world’s population suffered from allergic rhinitis, while over 300 million people were diagnosed with asthma^1^. However, the number of cases is increasing every year affecting patients’ quality of life, and ability to work, while having a significant impact on the global economy ^2,3^. Available allergy treatments are often described as inefficient or as causing side effects. Thus, it is necessary to search for alternative therapies^2^. In recent years, more studies have highlighted the role of healthy microbiota of the intestinal and airway tract in the prevention of allergy outcomes. Well-described are the differences between the microbiome composition of healthy and allergic patients^4^. Moreover, available studies indicate that the shape of the infant microbiome predicts allergic disease susceptibility in later life^5,6^. For instance, the Copenhagen Prospective Study on Asthma in Childhood (COPSAC) studied asthma susceptibility in infants and showed that hypopharynx colonization by pathogenic bacteria including *Streptococcus pneumoniae* or *Haemophilus influenzae* increased the risk of asthma development^7^. On the other hand, the nasal presence of *Lactobacillus* strains is associated with a lower risk of allergic asthma^4^. These breakthroughs have led to a new way of thinking, namely, introducing a health-beneficial bacteria to rebalance the microbiome and thus treat/prevent allergies. These microorganisms include bifidobacteria, which are anaerobic commensal bacteria well known for their immunomodulatory and health-promoting properties^8^. Also, they were proven to exhibit promising anti-allergic properties^9^. For instance, in an ovalbumin(OVA)-allergic mouse model, we have shown the preventive effect of *Bifidobacterium longum* ssp. *longum* CCM 7952 on the development of allergic sensitization and OVA-induced lung inflammation^10,11^. The additive effect of different bifidobacterial strains in reducing symptoms and improving quality of life in children with allergic rhinitis has also been shown in a clinical study^12^. Also, Ouwehand et al. (2009) described the combined influence of *Bifidobacterium lactis* Bl-04 and *Lactobacillus acidophilus* NCFM^TM^ on the alleviation of respiratory allergy symptoms and eosinophilic infiltration into nasal mucosa in children with birch pollen allergy ^13^.

Despite the promising results obtained for probiotics administration, several limitations remain. First, as the definition of probiotics claims, they are “live microorganisms that, when administered in adequate amounts, confer a health benefit on the host”^14^. Live organisms are more difficult to maintain and also their stability is low. Furthermore, they still possess the ability to reproduce, causing a risk of bacteriemia or the transfer of antibiotic-resistance genes^15,16^. Moreover, the complexity of a whole bacterial cell makes it impossible to determine its structure-function relationship in detail. For this reason, a new attempt has been proposed. So-called postbiotics started to gain attention in recent years and have been described by the International Association of Probiotics and Prebiotics as the “preparation of inanimate microorganisms and/or their components that confers a health benefit on the host”^15^. This definition includes different antigens present on the bacterial surface, such as proteins, glycolipids, lipoteichoic acids, and polysaccharides (PS).

We have recently described BAP1, the PS isolated from the surface of *Bifidobacterium adolescentis* CCDM 368, as a linear hexasaccharide with a unique structure consisting of glucose, galactose, and rhamnose residues, and a molecular mass of approximately 9.99 × 10^6^ ^17^. This polymer has been shown to be efficiently engulfed by airway epithelial cells and transferred to dendritic cells in *in vitro* assay. Treatment of bone marrow dendritic cells with the tested PS resulted in higher levels of TNF-α, IL-10, and IL-6. Further *ex vivo* immunomodulatory studies in OVA-restimulated splenocytes from OVA-sensitized BALB/c mice showed the ability of BAP1 to restore the balance between Th1/Th2-related cytokines, making it a very interesting molecule with anti-allergic potential.

Here, we first investigated the role of BAP1 in the naïve immune system development of germ-free (GF) mice, which, due to the sterile environment, were a perfect model for a precise study of the specific molecule-host reaction^18^. We further investigated the impact of BAP1 on the prevention and modulation of allergic (local and systemic) immune responses to OVA in a mouse model of allergy. The obtained results indicated the ability of BAP1 to alleviate allergic symptoms by re-establishing the Th1/Th2 balance.

## 2. Materials and methods

### 2.1. BAP1 PS

BAP1 is a surface PS from *Bifidobacterium adolescentis* CCDM 368 (Bad368) derived from human adult feces and made available for research by the Czech Collection of Dairy Microorganisms (CCDM, Laktoflora, Milcom, Tábor, Czech Republic). The bacteria cultivation and the PS isolation and purification were performed following the methods described previously^17^.

### 2.2. Animals

All animal procedures were performed following the EU Directive 2010/63/EU for animal experiments and were approved by the committee for the protection and use of experimental animals of the Institute of Microbiology, The Czech Academy of Sciences (no. 91/2019).

#### 2.2.1. GF mice

GF BALB/c mice (females, 3 weeks old) were kept under sterile conditions in Trexler-type plastic isolators and exposed to a 12 h light : 12 h dark cycle with unlimited access to autoclaved water and sterile irradiated diet (V1124-300, Ssniff Spezialdiäten GmbH, Germany).

#### 2.2.2. Specific-pathogen free mice

Specific-pathogen free (SPF) BALB/c mice (females, 6-8 weeks old) were kept in cages (IVC, Tecniplast, Italy) exposed to a 12 h light : 12 h dark cycle with unlimited access to water and sterile irradiated diet (V1124-300, Ssniff Spezialdiäten GmbH, Germany). SPF mice were regularly checked for the absence of potential pathogens according to an internationally established standard (FELASA).

### 2.3. Experimental designs

#### 2.3.1. GF mice treatment

GF mice (n = 6 per group) were divided into two experimental groups: PBS and BAP1. The experiment started with the intranasal (i.n.) administration of 30 μg/30 μl of BAP1 or PBS to 3-week-old mice anesthetized by isoflurane. PS administration was repeated an additional 2 times at one-week intervals (experimental design, Figure 1A). A week after the last treatment, mice were sacrificed and samples were collected.

**Figure 1.**
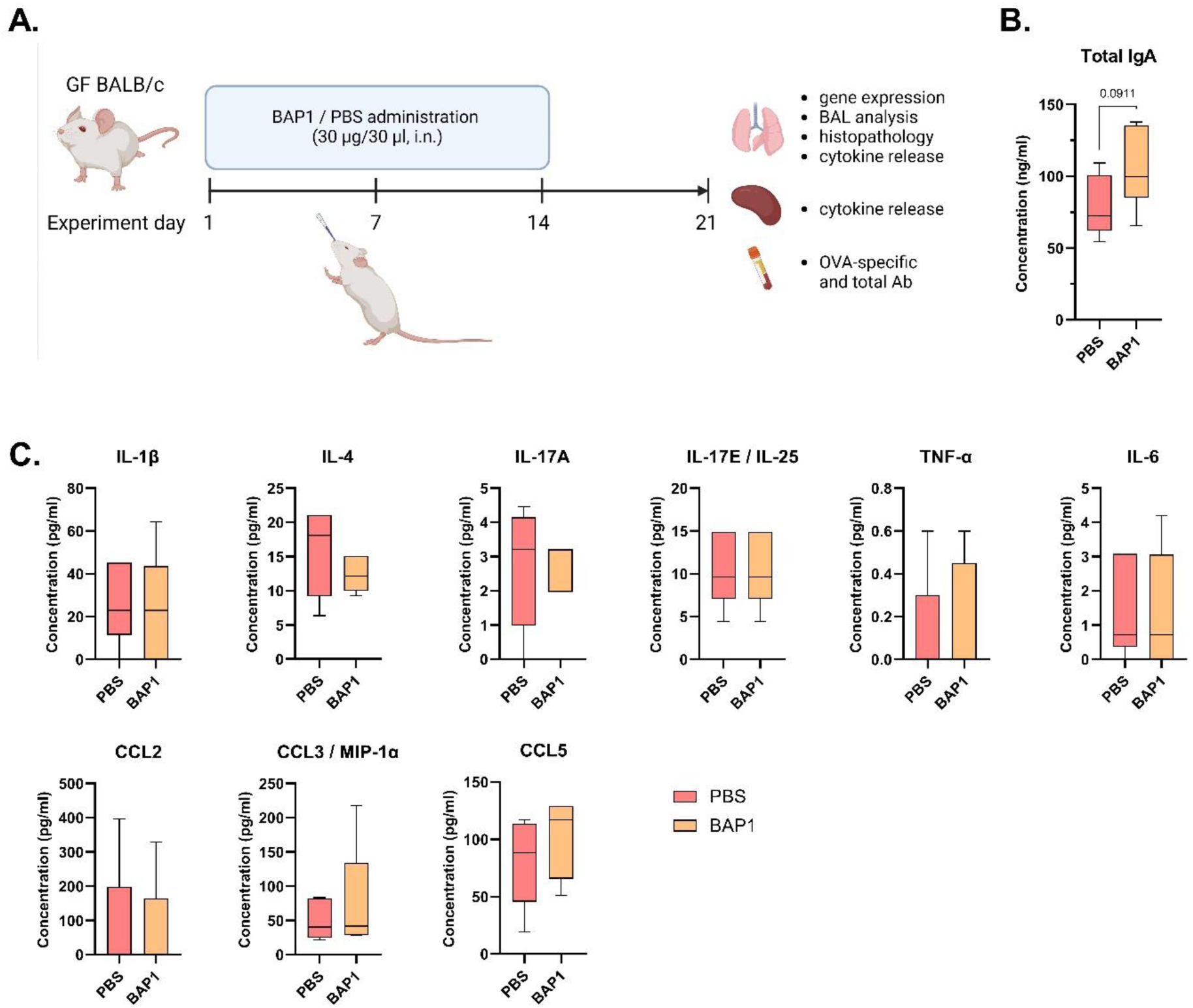
Systemic response of the GF mice to BAP1 i.n. application. **A.** Experimental design of GF mice treatment with PBS (control group) or BAP1 (30 μg / 30 μl / mouse). **B.** Total IgA serum antibodies tested by ELISA. **C.** Spontaneous cytokines and chemokines production in BAP1 and PBS-treated mice measured in splenocytes cultures by Luminex. An unpaired t-test was performed and significant differences between PBS- and BAP1-treated mice were calculated.

#### 2.3.2. Model of allergic airway inflammation to OVA

SPF mice (n = 5 – 6 per group) were divided into two experimental groups: OVA and BAP1. OVA-allergy model was introduced by 2 intraperitoneal (i.p.) injections of 10 μg of OVA (grade V, Sigma Aldrich, USA) in PBS (100 μl) mixed with Alum (100 μl; Alu-gel-S, SERVA Electrophoresis, GmbH, Germany) in a 14-day interval. Third boosting immunization was performed with 15 μg of OVA (grade V, Sigma Aldrich) in PBS (100 μl) mixed with Alum (100 μl) one week after the second injection. 4 hours before each i.p. sensitization, mice were treated intranasally with 30 μl of BAP1 (30 μg per dose) or PBS (OVA group). 7 days after the third immunization, 30 μl of PBS (OVA group) or BAP1 (30 μg /30 μl) were applied intranasally 4 hours before the OVA challenge (100 μg per dose, 30μl, intranasally) and repeated for 3 consecutive days. The following day, mice were sacrificed and samples were collected (experimental design, Figure 4A).

### 2.4. Splenocyte isolation

Splenocyte isolation was performed on aseptically removed spleens according to the method described by Pyclik et al. ^11^. Isolated cells were counted and seeded on a 96-well plate (1 × 10^7^ cells/ml, 100 μl/well) in RPMI 1640 medium (Sigma Aldrich) supplemented with 10 % FBS (Fetal Bovine Serum, Gibco), 100 U/ml of penicillin, 100 μg/ml streptomycin, and 10 mM HEPES (Sigma Aldrich). Splenocytes derived from GF mice were incubated without additional treatment, with a total volume of 200 μl of medium per well. Splenocytes isolated from the OVA-sensitized mice were restimulated on a plate with 100 μg of OVA (100 μl/well in a total volume of 200 μl). Finally, cells were incubated for 72 h at 37 °C (5 % CO_2_, appropriate humidity). The concentration of cytokines was measured in supernatants by the Milliplex Map Mouse Cytokine/Chemokine Panel including CCL2, CCL5, CCL3, IL-6, IL-1β, TNF-α, IL-17E, IL-4 for GF mice and IL-10, IFN-γ, IL-4, IL-5, and IL-13 for OVA-allergy model. The procedure was performed according to the manufacturer’s instructions and analyzed with the Luminex 2000 System (Bio-Rad Laboratories, USA). Unstimulated cells were used as a background for restimulated cells.

### 2.5. Lung cells

The left lung lobe was aseptically collected from BALB/c mice, and placed in a digestion buffer containing Liberase TL (0.05 mg/ml, Sigma Aldrich) and DNAse (0.5 mg/ml, Sigma Aldrich) dissolved in RPMI 1640 medium supplemented with 100 U/ml of penicillin, 100 μg/ml streptomycin, and 2 mM L-glutamine. To increase the efficiency of the process, the lungs were cut into smaller pieces with sterile scissors and left for 45 min at 37 °C. Digested cells were pressed through a 70 μm cell strainer and centrifuged (1300 rpm, 10 min, 4 °C). Pellet was treated with ACK lysis buffer for 3 min and the reaction was stopped with supplemented RPMI 1640 medium completed with 10 % FBS. Finally, centrifuged cells were counted and seeded on a 96-well plate (0.2 × 10^6^ cells/well).

Cells derived from GF mice were incubated without additional treatment, with a total volume of 200 μl of medium per well. Cells from the OVA-allergy model were restimulated on a plate by 100 μg of OVA (100 μl/well in a total volume of 200 μl). Finally, plates were incubated for 72 h at 37 °C (5 % CO_2_, appropriate humidity). The concentration of cytokines was measured in supernatants by the Milliplex Map Mouse Cytokine/Chemokine Panel including CCL2, CCL5, IP-10, CCL3, CCL11, IL-6, IL-1β, TNF-α, IL-17A, IL-17E, IL-4 for GF mice and IL-10, IFN-γ, IL-4, IL-5, and IL-13 for OVA-allergy model. The procedure was performed according to the manufacturer’s instructions and analyzed with the Luminex 2000 System (Bio-Rad Laboratories). Unstimulated cells were used as a background for restimulated cells.

### 2.6. Histopathological evaluation of lungs

The middle lobe of the lungs collected from the GF and OVA-allergy mouse model was fixed in 4 % paraformaldehyde for 24 h and stored in 80 % ethanol. Fixed lungs were embedded in paraffin, cut into 5 µm-thick slides, and subjected to periodic acid-Schiff staining (PAS). Prepared sections were assessed for a histological score with light microscopy (100 × magnification) as described by Srutkova et al. ^18^. Samples were viewed under Olympus BX 40 microscope equipped with an Olympus Camedia DP 70 digital camera, and the images were analyzed using Olympus DP-Soft. The average histological score was assessed in 5 fields of the slide per each mouse and expressed as the sum of single parameters (perivascular and peribronchiolar inflammation, number of cells in alveolar spaces, and a number of PAS-positive cells) described in Table 1 and divided by 3.

**Table 1.**
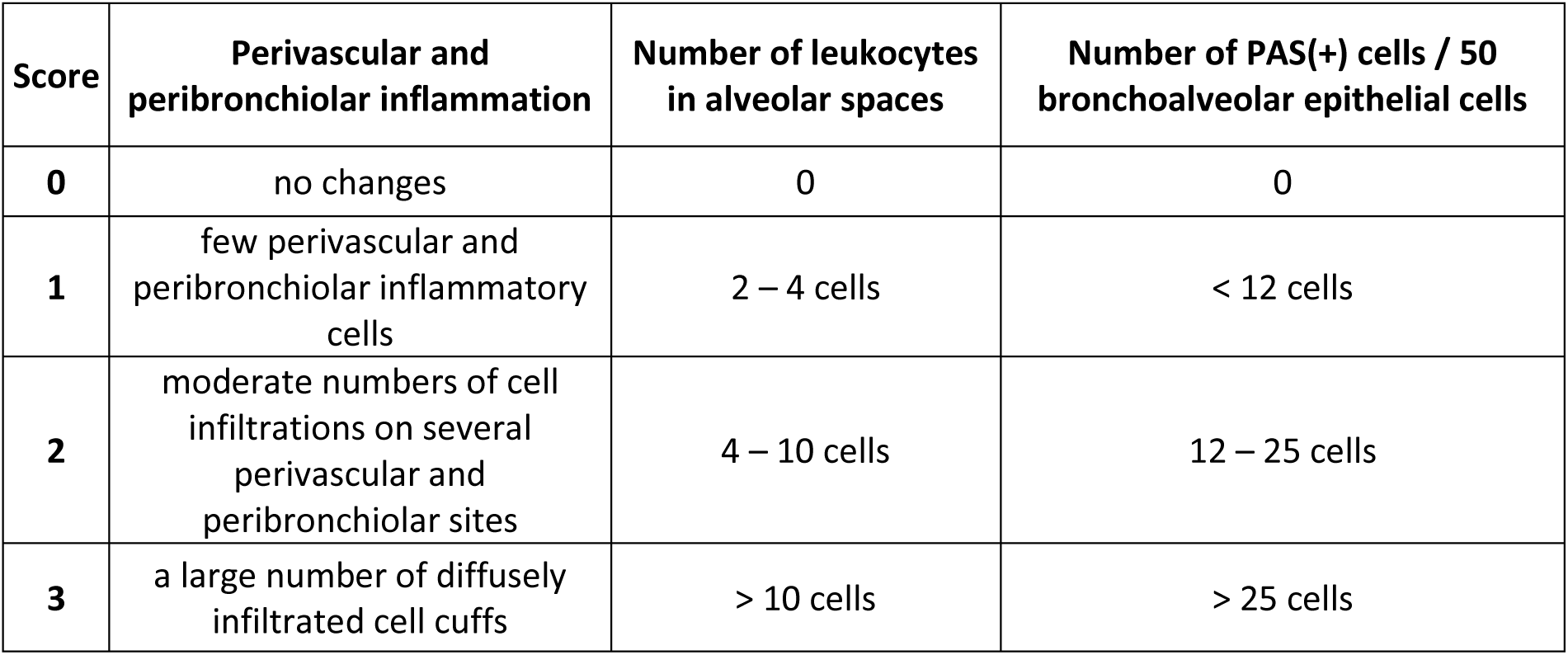
Principles of histological sections assessment.

### 2.7. Investigation of bronchoalveolar lavage (BAL)

To collect bronchoalveolar lavage fluid (BALF), the lungs were gently flushed with 2 × 0.5 ml of sterile Dulbecco’s Phosphate Buffered Saline (D-PBS, Gibco) through trachea cannulation. The obtained suspension was centrifuged (400 × g, 7 min, 4°C) and the supernatant was separated from the pellet and stored at – 80 °C. The cell pellet was resuspended in 200 µl of RPMI 1640 medium (Sigma Aldrich) and used for cell differential count. Briefly, BAL cells were transferred to a microscope slide with a cytospin, left to dry, fixed with methanol, and stained with a Dip-Quick-stain kit following the manufacturer’s instructions (Medical Products, Czech Republic). BAL cells were evaluated with light microscopy following standard morphologic criteria (200 cells observed per cytospin preparation) ^19^.

### 2.8. Immunological evaluation of sera and BALF

Total IgA, IgG-Fc, and IgE antibody levels were detected with Bethyl kit (Bethyl Laboratories, Inc), according to the manufacturer’s recommendations. Briefly, goat anti-mouse antibodies were used to coat the 96-well microtiter plates (Nunc MaxiSorp, Thermo Fisher Scientific), wells were blocked with 1 % bovine serum albumin (BSA), and tested sera or BALFs were added. After rinsing the excess reagents, HRP-conjugated detection antibodies were incubated on the plate for 1 h, and TMB substrate was used to develop a colorimetric reaction. Finally, the reaction was stopped with 2 M H_2_SO_4_ and the absorbance was read at 450 and 570 nm.

Antigen-specific IgA, IgE, IgG-Fc, IgG1, and IgG2a antibody levels were detected with Bethyl reagents (Bethyl Laboratories, Inc), according to the manufacturer’s recommendations. Briefly, 0.5 μg/well of OVA in PBS was used to coat the 96-well microtiter plates, wells were blocked with 1 % BSA in PBS, and diluted sera or BALFs were added. HRP-conjugated anti-mouse detection antibodies were used and TMB substrate was used. The reaction was stopped by adding 2 M H_2_SO_4_ and the results were assessed by an absorbance read at 450 and 570 nm.

### 2.9. RT-qPCR

RNA was isolated from the superior lobe of the lung with a NucleoSpin® RNA kit (Macherey-Nagel) and cDNA was prepared with the use of SuperScript™ II Reverse Transcriptase (Invitrogen) kit according to the manufacturer’s instructions. For Real-Time PCR, a master mix was prepared for each of the tested genes including *Actb* – reference gene, *Rorc*, *Gata3*, *Tbx21*, *Foxp3*, and *Il10*, that included (for 1 well): 5 μl of the SYBR Green Real-Time PCR master mix (Promega), 1.25 μl of the forward and reverse primer (standardized KiCqStart^TM^ primers, Merck). 7.5 μl of the prepared mixes were added to white 96-well qPCR plate wells with 2.5μl of the tested cDNA. The mRNA expression was calculated with the 2–ΔΔCT. Analysis was performed with CFX Connect^TM^ Real Time System equipped with Thermal Cycler (Serial No. BR002307; BioRad) and Optical Module (Serial No. 788BR02394; BioRad). Gene expression was counted in relation to the mean of negative control mice (PBS-treated GF and OVA-treated mice).

### 2.10. Statistical analysis

All data are presented as boxes and whiskers with median + minimal and maximal values. An unpaired Student’s t-test was used to compare results between PBS- and BAP1-treated GF mice and between PBS- and BAP1-treated OVA-sensitized and challenged mice. All statistically significant results were marked on the graphs (∗∗∗∗p < 0.0001, ∗∗∗p < 0.001, ∗∗p < 0.01, ∗p < 0.05). GraphPad Prism 10 Software (San Diego, CA, USA) was used for statistical calculations and visualizations.

## 3. Results

### 3.1. BAP1 treatment increases IgA levels in GF mice

To evaluate the impact of BAP1 on the naïve immune system, we tested it in GF mice. PS was administered to mice on the experimental day 1, 7, and 14. On the 21^st^ day, mice were sacrificed and spleen, lungs, sera, and BALF were collected (Figure 1A). To assess the systemic responses, the levels of total IgA, IgG-Fc, and IgE antibodies were measured in sera. Results showed a trend to increase total IgA compared to mice treated with PBS only (Figure 1B, Supplementary Figure 1). We examined the BAP1-specific recall response in the spleen and observed that there were no significant changes in spontaneous cytokine/chemokine production when compared to the control group (Figure 1C).

### 3.2. BAP1 administration does not induce inflammatory lung responses in GF mice but activates Rorc expression

Further evaluation of the BAP1 influence on the lung immune responses of GF mice showed no differences in the total cell or different immune cell types (such as macrophages or lymphocytes) count in the BAL (Figure 2A). Importantly, we did not observe an undesirable infiltration of inflammatory neutrophils or eosinophils to the lung tissue. Simultaneously, we detected a significant increase in the total IgA antibody levels in BALF (Figure 2B). Histopathological examination of the lung tissue confirmed the absence of aberrant immune cell infiltration or other morphological changes in response to BAP1 treatment (Figure 2C). In the lung cell culture, administration of BAP1 to GF mice was associated with a significant decrease in IL-6 and CCL-2 levels and a tendency to inhibit eotaxin production. On the other hand, BAP1 administration increased the spontaneous CCL5 production (Figure 2D). To investigate the impact of BAP1 on the activation of certain transcription factors, the upper lobe of the lung was subjected to RNA isolation. We examined four transcription factors *Gata3*, *Tbx21*, *Foxp3*, and *Rorc*, which are involved in Th1, Th2, Treg, and Th17 responses respectively. Our results indicated that PS treatment significantly induced the *Rorc* gene and showed a tendency to increase *Tbx21* expression in comparison to the PBS control group (Figure 2E).

**Figure 2.**
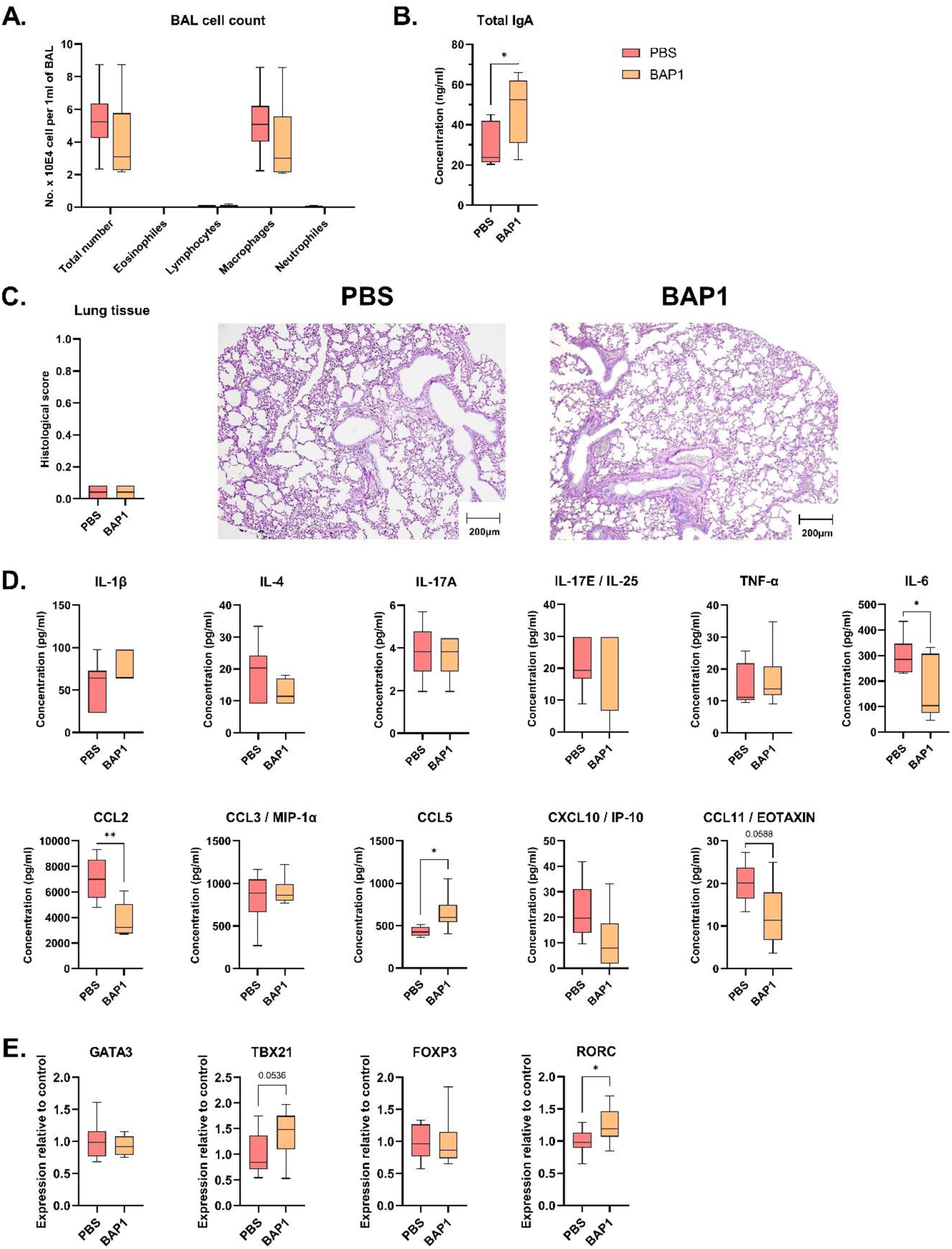
Lung-related response of GF mice to BAP1 treatment. **A.** Cell count in BAL. **B.** Total IgA BALF antibodies tested by ELISA. **C.** Histopathological analysis and representative histopathological section of lungs from mice treated with PBS or BAP1 stained with Periodic Acid-Schiff (magnification 100x, scale 200 um). **D.** Spontaneous cytokines and chemokines production in PBS- and BAP1-treated mice measured in lung cell cultures by Luminex. **E.** Gene expression detected in lungs of GF mice treated with PBS or BAP1. An unpaired t-test was performed and significant differences between PBS- and BAP1-treated mice were calculated (* p ≤ 0.05).

### 3.4. BAP1 treatment reduces OVA-specific IgE levels in mice with allergic airway inflammation

Given the established immunomodulatory potential of BAP1, we tested the ability of BAP1 to alleviate allergy symptoms in a mouse model of allergic airway inflammation. Experimental design was performed according to our previous study with 3 i.p. OVA-injections and 4 i.n. challenges. BAP1 was administered intranasally to mice 4 h prior to each OVA sensitization and challenge (Figure 3A) ^11^. The levels of total IgA and IgE antibodies in the sera were not notably affected by BAP1 treatment, but we observed a trend towards a reduction in the OVA-specific IgA production in sera. Moreover, PS administration dampened Th2 systemic humoral sensitization, as the OVA-specific IgE level was significantly reduced. We observed a significant increase in the production of OVA-specific IgG1 antibodies in serum and no changes in OVA-specific IgG2a (Figure 3B, Supplementary Figure 2). At the same time, a significant reduction in the level of Th2-related IL-5 cytokine, and a tendency to smaller production of IL-10, IL-13, and IFN-γ was detected in splenocytes stimulated by OVA (Figure 3C).

**Figure 3.**
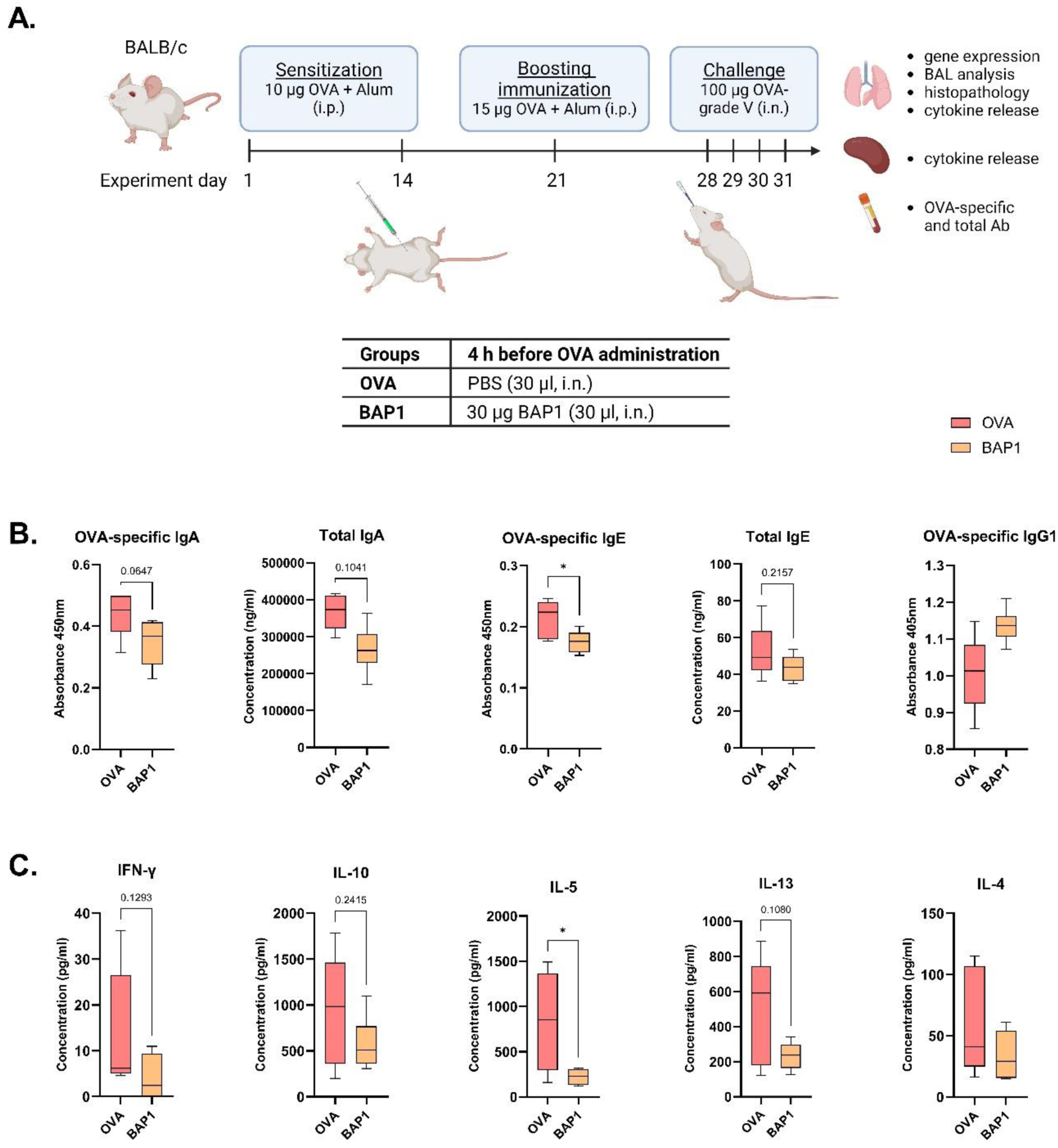
Systemic response after i.n. administration of BAP1 in the mouse model of allergic airway inflammation. **A.** Experimental design for BAP1 treatment, OVA sensitization and challenge. **B.** OVA-specific and total IgA, IgE, and IgG serum antibodies tested by ELISA. **C.** Cytokines and chemokines production in PBS- and BAP1-treated mice in response to restimulation with an allergen (OVA) measured in splenocyte cultures by Luminex. An unpaired t-test was performed and significant differences between PBS- and BAP1-treated mice were calculated (* p ≤ 0.05).

### 3.5. BAP1 reduces airway inflammation and decreases the expression of the Il10 gene

Directly in the lung tissue, i.n. BAP1 treatment was able to reduce eosinophil and macrophage numbers without affecting lymphocyte or neutrophil cell populations (Figure 4A). Evaluation of the total and OVA-specific IgA levels in BALF showed no differences between the studied groups (Figure 4B).

**Figure 4.**
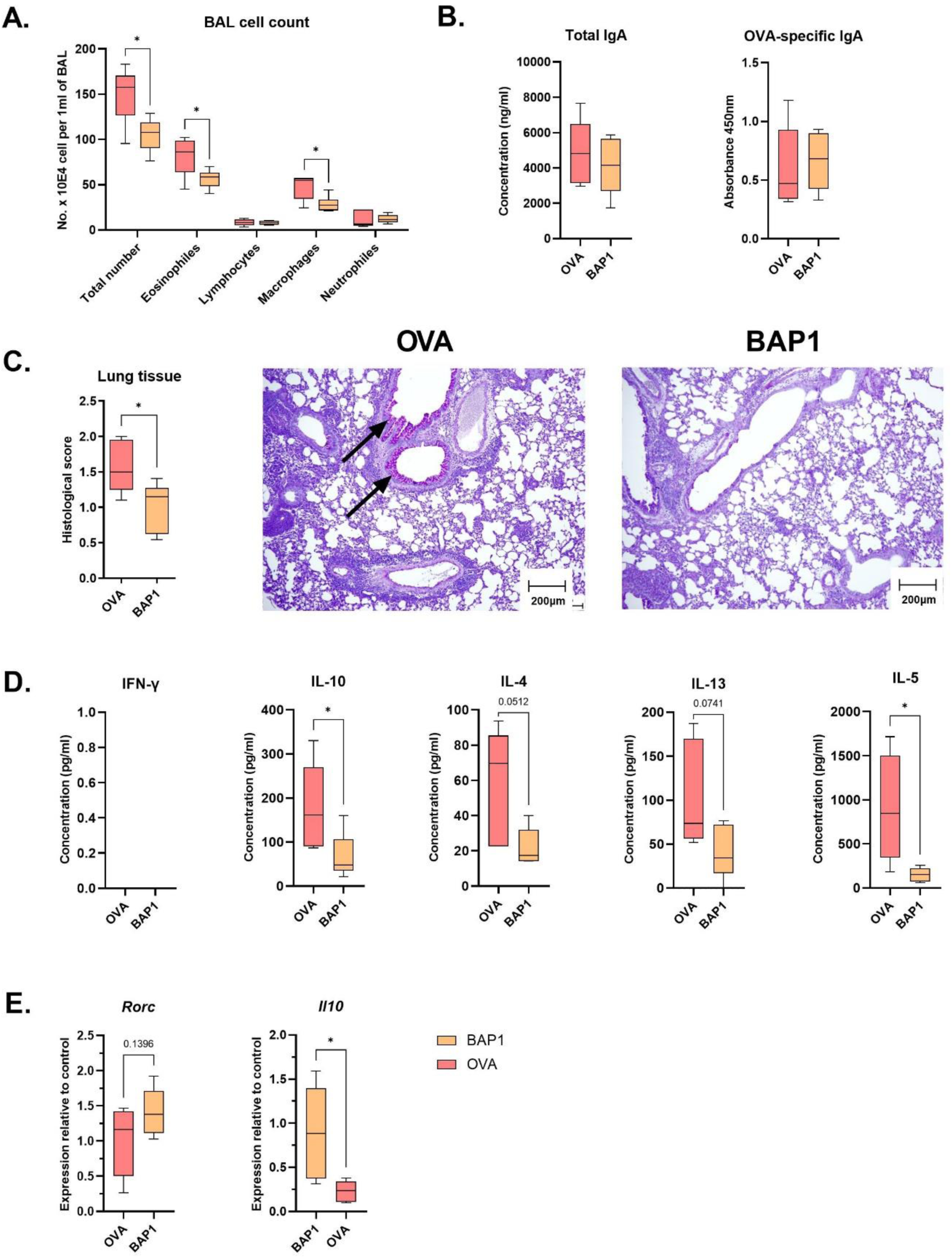
Lung-related response to BAP1 in mice with allergic airway inflammation. **A.**Cell count in BAL. **B.** Total and OVA-specific IgA BALF antibodies tested by ELISA. **C.** Representative histopathological section of lungs from OVA-sensitized mice treated with PBS or BAP1, stained with Periodic Acid-Schiff. **D.** Cytokines and chemokines production of BAP1-treated and OVA-control mice restimulated with an allergen (OVA) measured in lung cell cultures by Luminex and expressed as pg/ml. **E.** Changes in lung gene expression measured in BAP1- and PBS-treated OVA-allergy mice. An unpaired t-test was performed and significant differences between OVA-allergy mice treated with PBS and BAP1 were calculated (* p ≤ 0.05, ** p ≤ 0.01, *** p ≤ 0.001).

However, histological analysis of the lung tissue showed a reduction in leukocyte infiltration in the perivascular, peribronchiolar, and alveolar spaces, and a reduction in mucus-producing cells in the airways of BAP1-treated mice (summarized in the histopathological evaluation in Figure 4C). Similar to the cytokine recall response in splenocyte cultures, BAP1 treatment significantly downregulated the OVA-induced IL-5 and IL-10 levels and decreased the IL-4 and IL-13 production in lung cell cultures (Figure 4D).

Evaluation of lung gene expression showed a tendency to increase *Rorc* levels, which is consistent with our findings in GF mice (Figure 4E). The analysis revealed a significant inhibition of *Il10* gene expression (Figure 4E). In addition, investigation of the remaining genes, including *Gata3*, *Tbx21*, and *Foxp3,* showed no changes between tested groups of mice (Supplementary Figure 3).

## Discussion

In our recent study, we evaluated the role of surface molecules isolated from the *Bifidobacterium adolescentis* CCDM 368 strain using *in vitro* and *ex vivo* studies ^17^. We showed that surface PS, BAP1, is an immunoregulatory molecule that is well-recognized and transferred by epithelial and immune cells. Its ability to restore the Th1/Th2 balance (disturbed in allergy) was highly desirable, as demonstrated by the induction of IFN-γ and the inhibition of IL-13 and IL-5 production in OVA-sensitized splenocyte cultures. However, to fully understand the exact nature of the molecule, it is necessary to know not only its function but also its structure. Thus, we determined the repeating unit of the BAP1, which turned out to be a unique, non-branched PS consisting of glucose, rhamnose, and galactose residues with a molecular mass of approximately 9.99 × 10^6^ ^17^.

Here, to expand our knowledge about the BAP1 role in the immune response modulation, we tested it in a naïve system of airways in GF mice to observe the molecule-specific responses (Figure 1A). We noticed an increase in total IgA serum levels in BAP1-treated GF mice which is consistent with the previously published data (Figure 1B)^20,21^. Enhanced levels of this immunoglobulin are crucial for balanced microbial colonization and induction of immune tolerance against introduced microbiota or beneficial bacteria-derived antigens ^18,22^. Further evaluation of splenocyte response showed no differences in cytokine and chemokine production after treatment of GF mice with BAP1 (Figure 1C). The neutral effect of tested PS favors our work, since it indicates, that BAP1 doesn’t cause a systemic inflammatory response of the naïve immune system.

For the next step, we decided to focus on the immune response directly in the lungs of GF mice, as it is one of the main organs affected by airway allergy inflammation. Evaluation of cell count in BALF showed no differences between the PS-treated and control groups, however, the total IgA level was significantly increased (Figure 2A and B). Similarly, increased levels of total IgA in GF mice lungs have been shown upon bacterial PS administration in our previous study ^18^. Also, the PAS staining indicated no evidence of inflammatory signs in the lung tissue (Figure 2C). Interestingly, we observed a significant decrease in CCL2 (C-C Motif Chemokine Ligand 2) protein levels in lung cells supernatants (Figure 2D). This chemokine is known to be upregulated in airway allergy, leading to the recruitment of macrophages and basophils, thereby, inducing the development of inflammation ^23^. Activation of CCL2 is mediated via Th2 cytokines, indicating, that our PS may have an inhibitory effect on the production of cytokines such as IL-4 ^24^. We also observed a trend towards decreased levels of CCL11/eotaxin, which recruits eosinophils, key factors in type 2 eosinophilic asthma (Figure 2D) ^25^. Finally, we complemented our analysis in GF mice by investigating the expression of transcription factors *Tbx21*, *Gata3*, *Foxp3*, and *Rorc*, which are responsible for the activation of four different pathways – Th1, Th2, Treg, and Th17 respectively. The results pointed to the possible role of BAP1 in activating the Th17 and Th1 pathways, as we observed an increased level of *Rorc* gene expression and a tendency to increase the *Tbx21*, which in turn could restore the balance of the immune responses in allergic diseases (Figure 2E).

Considering the data obtained in GF mice, and the previously described *ex vivo* anti-allergic potential, we continued the BAP1 studies *in vivo* using the mouse model of allergic airway inflammation to OVA. The available research describing the impact of beneficial bacteria on the development of respiratory allergic diseases has focused mainly on the oral administration of whole bacteria. For instance, Cavalcanti et al. (2024) described the anti-allergy potential of *Lacticaseibacillus paracasei* on the OVA-induced inflammation in BALB/c mice ^26^. Results showed an inhibitory effect of the investigated bacteria to reduce nasal and lung tissue inflammation and decrease BALF eosinophils number and OVA-specific IgE in sera. Also, the strain mitigated Th2-related cytokines and TGF-β production, while increasing IL-10 levels. A similar effect in alleviating airway allergy was observed for other orally delivered lactic acid bacteria strains ^27–31^. However, only a few publications focus on the i.n. administration of bacteria or bacterial antigens ^11,18,32,33^. This attempt allows to achieve a stronger local response to the treatment. One of the advantages of i.n. probiotic administration is that bacteria do not pass through the digestive system, and thus are not exposed to the digestive enzymes. Spacova et al. investigated the i.n. application of *Lactobacillus rhamnosus* GG to birch pollen-induced BALB/cOlaHsd. The results showed that bacteria treatment reduced inflammation and eosinophil infiltration in the lungs and inhibited Th2-related IL-5, IL-13, and IL-10 production ^33^. Previously, we tested i.n. administration of EPS from *Lacticaseibacillus rhamnosus* LOCK900 to OVA-allergy mice and observed the alleviation of allergy symptoms through inhibition of Th2-related cytokine responses and decreased number of eosinophils ^34^. Therefore, based on available data and our own experience, in this study, we focused on i.n. administration of the BAP1 molecule as the most effective route to modulate the immune response in a mouse model of airway allergy inflammation.

Next, we evaluated the systemic response to BAP1 treatment. Investigation of serum levels of total and allergen-specific IgA and IgE showed the ability of BAP1 to inhibit the production of both immunoglobulins with the significant reduction of OVA-specific IgE (Figure 3B). We observed a similar trend in our previous studies regarding L900/2 and L900/3 PSs isolated from *Lactobacillus rhamnosus* LOCK 0900, where tested PS significantly reduced allergen-specific IgE and showed a tendency to mitigate specific IgA levels ^34,35^. Investigation of splenocyte culture restimulated with OVA showed a role of BAP1 in the inhibition of the Th2-related cytokines, responsible for the development of allergy inflammation and the recruitment of IgE antibodies. That included a significant inhibition of IL-5 and a tendency to decrease IL-4 and IL-13 cytokines (Figure 3C).

Research focusing on the impact of probiotic bacteria on the development of allergies indicates the beneficial effect of lactic acid bacteria in reducing cell infiltration thus alleviating cell inflammation and tissue damage ^11,27,30,31^. Further investigation in the lung has yielded promising results in favor of the anti-allergic potential of BAP1. Histopathology evaluation of the lung tissue showed significantly reduced inflammation in BAP1-treated mice compared to OVA mice, accompanied by a reduced number of cells counted in BALF. Particularly, we saw a decrease in macrophages as well as eosinophils, crucial for IgE-mediated inflammation (Figure 4A and C). Reduced eosinophilia upon probiotic-treatment is well described in the literature and is associated with anti-allergic properties of the strain in the airway tract ^36,37^. Also, postbiotic impact on cell infiltration was described in the literature. PS isolated from *Bifidobacterium longum* subsp. *longum* 35624™ significantly inhibited eosinophil infiltration in BAL of OVA-sensitized mice ^38^.

Further, BAP1 treatment downregulated the Th2 allergic cytokine response locally in lung cells and systemically in splenocytes (Figure 4B and D). Generally, the available research mostly agrees that probiotics’ mechanism of restoring disrupted Th1/Th2 balance is based on the increased production of IFN-γ or IL-10 ^39^. Interestingly, in both systemic and local responses, the level of IL-10 was significantly lower upon BAP1 treatment in comparison to OVA mice. At first, studies focusing on IL-10 associated this cytokine only with Th1/Th2 functionalities. However, current knowledge underlines the complexity of this molecule and shows a broad range of cells that can produce IL-10 for different anti-, pro-inflammatory, or regulatory purposes ^40,41^. A great number of studies show the protective role of IL-10 in the development of asthma diseases and the role of probiotics in increasing this cytokine. ^26,30,31,42^. However, there is evidence that it can also be responsible for increasing airway inflammation. Polukort et al. described IL-10-dependent mast cell activation affecting the IgE-mediated food allergy development in OVA-sensitized BALB/c mice ^43^. Wu et al., 2016 have shown that administration of *L. rhamnosus* GG before or after the induction of OVA inflammation reduced allergy symptoms while decreasing IL-10 cytokine levels, in both serum and BALF. This proves that the alleviation of allergy can be associated with mitigated levels of IL-10 ^27^. Interestingly, examination of *Rorc* gene indicated an increased expression, as in GF mice administered with BAP1, however, the effect was statistically insignificant (Figure 4E). The proper control of the *Rorc* gene is crucial for the prevention of allergy development. Abdel-Gadir et al., described the influence of Clostridiales species on food allergy in mice. In addition to describing the bacterial suppressive effect on allergy inflammation, they investigated its signaling. The results highlighted the role of ROR-γ Treg cells in disease alleviation ^44^. The described process is a turnover in current knowledge, since, until now, the ROR-γ molecule was mainly associated with the allergy occurrence. We are aware that the mechanism differs between food and airway hyperresponsiveness, but this example shows that there’s still much to be discovered about the mechanisms that underlie the beneficial effect of bacteria in alleviating allergies.

It is worth noticing that in our studies, we decided to go further and evaluate the effect of bacterial components instead of the whole microorganism. Through this attempt, we could overcome the difficulties connected to the use of a live microorganism such as the transfer of antibiotic resistance genes or bacteriemia ^15,45^. Moreover, PSs are heat resistant, thus more stable molecules, and in comparison to the bacteria cells, it is possible to define their exact structure and function ^46^. Only a few publications focus on structure-function studies of PSs ^34,47–49^. Even less discuss PSs properties to treat allergies. Bai et al. described LJP heterogenous PS from *Lonicera japonica* that when administered intragastrically was able to reduce symptoms of OVA-induced allergy in mice by inhibition of serum IgE levels, nasal eosinophils number, allergy-related cytokines, and expression of *Rorγt* genes in the nasal mucosa ^49^. Also, in our previous work, we confirmed an inhibitory effect of exopolysaccharide (EPS) of *Lacticaseibacillus rhamnosus* LOCK900 on the development of allergy inflammation in OVA-treated mice ^34^. This effect was assessed by i.n. administration of PS, similar to BAP1. However, the structure of the molecule and possible mechanisms involving IgA and TGF-β activation are distinct from the one presented in this publication.

## Conclusions

Here, for the first time, we conducted a detailed analysis of BAP1 PS to evaluate its anti-allergic potential in a mouse model of OVA allergy. Primarily, our results confirmed the neutral effect of BAP1 on the naïve immune system of GF mice. Nevertheless, we observed a significant CCL2 downregulation, a decrease in eotaxin production, and an upregulation of *Rorc* gene expression in lung. Further studies performed on OVA-treated mice showed the ability of BAP1 to reduce allergy inflammation. This effect was achieved by inhibition of OVA-specific IgE and Th2-related cytokine production as well as eosinophil and macrophage recruitment. Moreover, treatment with BAP1 inhibited the level of Th2-related and IL-10 cytokines. Finally, we were able to indicate the possible role of increased *Rorc* and decreased *Il10* gene expression in the BAP1 mechanism. In conclusion, BAP1 appears to be a promising candidate for the management of allergic diseases. Nevertheless, modulation of immune response to bacterial antigens is complex, and many factors, such as cells and tissue type, genetic and environment immunity predisposition, microbiome, etc. need to be taken into account. Also, there is little information about possible PSs’ molecular pathways. Thus, further studies will be required to fully confirm the mechanism and potential of BAP1.

## CRediT authorship contribution statement

**Katarzyna Pacyga-Prus:** Methodology, Formal analysis, Investigation, Resources, Writing - original draft, Visualization. **Tereza Hornikova:** Investigation, Formal analysis, Writing - review & editing. **Dagmar Srutkova:** Methodology, Resources, Writing - review & editing, Funding acquisition. **Katarzyna Leszczyńska-Nowak:** Investigation**. Agnieszka Zabłocka**: Writing - review & editing **Martin Schwarzer:** Resources, Writing - review & editing, Supervision, Funding acquisition. **Sabina Górska:** Conceptualization, Validation, Writing - review & editing, Supervision, Project administration, Funding acquisition.

## Acknowledgments

We thank Jaroslava Valterova and Sarka Maisnerova for their excellent technical assistance. Graphical abstract and experimental models were prepared using Biorender online software.

