## Supplementary Materials for "Polysaccharide BAP1 of *Bifidobacterium adolescentis* CCDM 368 attenuates ovalbumin-induced allergy through inhibition of Th2 immunity in mice"

**Suplementarny materials**

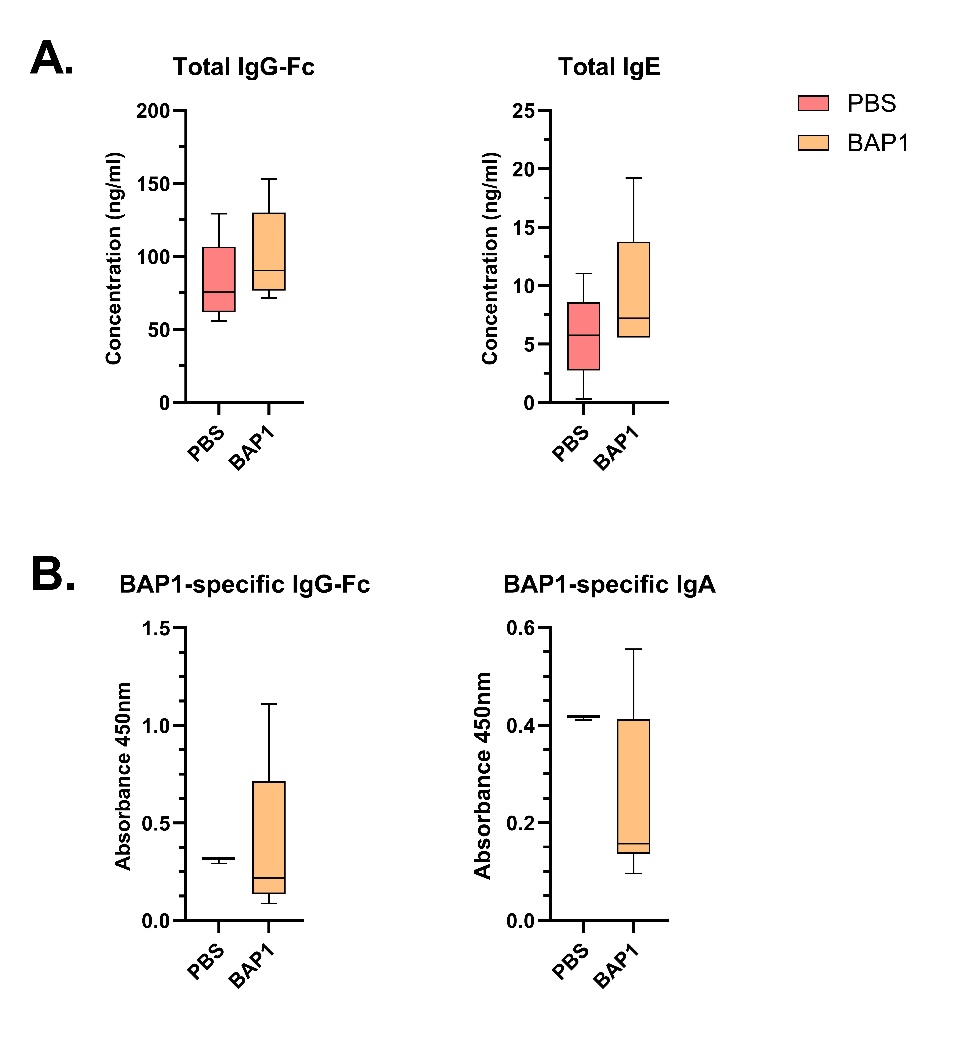

**Supplementary Figure 1.** Total IgG-Fc and IgE, serum antibodies measured in GF mice and tested by ELISA. An unpaired t-test was performed and significant differences between PBS and BAP-1 treated mice were calculated.

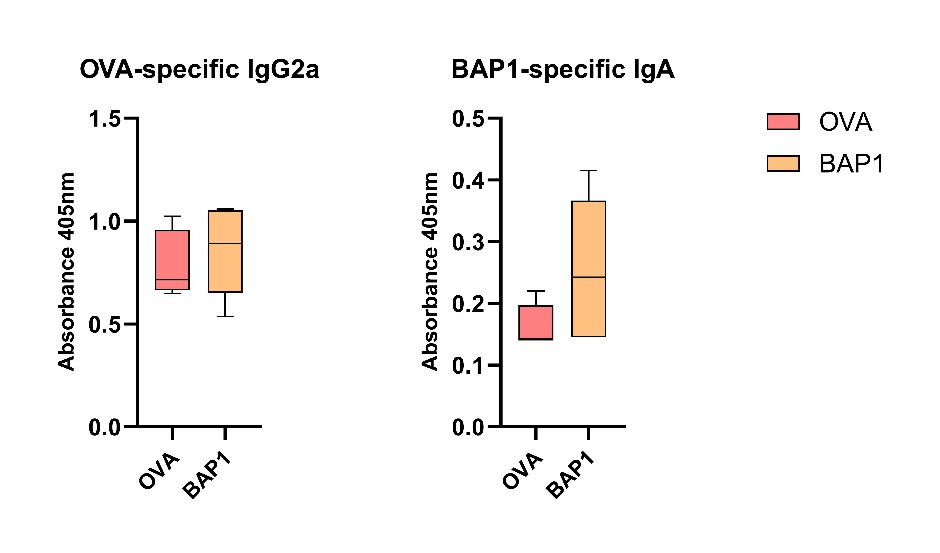

**Supplementary Figure 2.** OVA-specific IgG2a serum antibodies measured in OVA-induced allergic mice and tested by ELISA. An unpaired t-test was performed and significant differences between PBS and BAP-1 treated mice were calculated (* p ≤ 0.05).

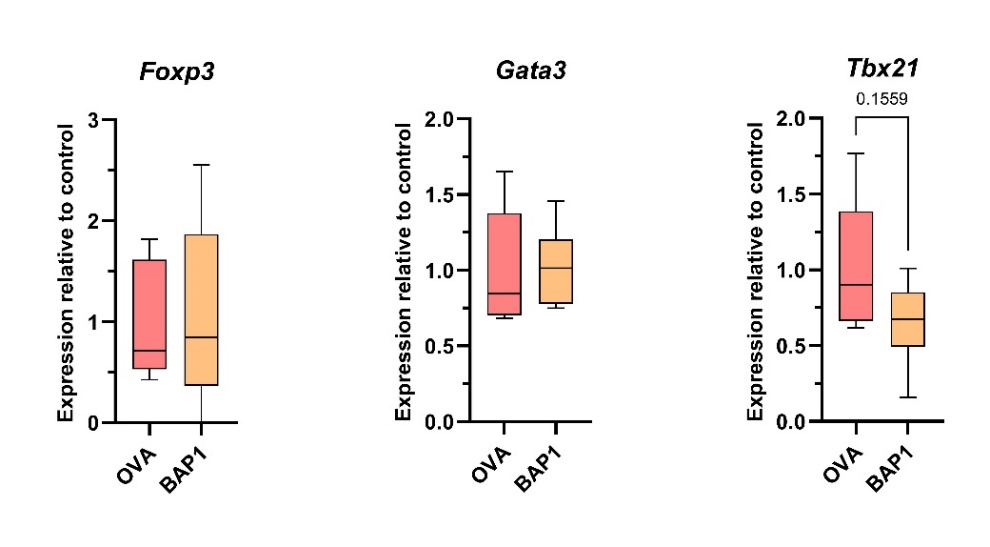

**Supplementary Figure 3.** Changes in lung gene expression measured in BAP-1 treated OVA-allergy mice. An unpaired t-test was performed and significant differences between PBS and BAP-1 treated mice were calculated.
